# Benchmarking gene embeddings from sequence, expression, network, and text models for functional prediction tasks

**DOI:** 10.1101/2025.01.29.635607

**Authors:** Jeffrey Zhong, Lechuan Li, Ruth Dannenfelser, Vicky Yao

## Abstract

Accurate, data-driven representations of genes are critical for interpreting high-throughput biological data, yet no consensus exists on the most effective embedding strategy for common functional prediction tasks. Here, we present a systematic comparison of 38 gene embedding methods derived from amino acid sequences, gene expression profiles, protein–protein interaction networks, and biomedical literature. We benchmark each approach across three classes of tasks: predicting individual gene attributes, characterizing paired gene interactions, and assessing gene set relationships while trying to control for data leakage. Overall, we find that literature-based embeddings deliver superior performance across prediction tasks, sequence-based models excel in genetic interaction predictions, and expression-derived representations are well-suited for disease-related associations. Interestingly, network embeddings achieve similar performance to literature-based embeddings on most tasks despite using significantly smaller training sets. The type of training data has a greater influence on performance than the specific embedding construction method, with embedding dimensionality having only minimal impact. Our benchmarks clarify the strengths and limitations of current gene embeddings, providing practical guidance for selecting representations for downstream biological applications.

## Main

Gene embeddings have been increasingly recognized as powerful tools in computational biology, providing a framework to transform complex, high-dimensional biological data into compact, manageable numerical vector representations. Embeddings serve as a bridge between raw biological datasets and machine learning models, allowing researchers to efficiently extract meaningful patterns that may be otherwise obscured by noise and heterogeneity in the data. By amplifying critical biological signals, embeddings can significantly improve the performance of downstream predictive tasks, including disease associations [1, 2], gene function [3–5], and genetic interactions [6], among many other tasks in the biomedical realm.

In recent years, a variety of gene embeddings have emerged [7], expanding on classic biological vector encodings [8–13], by unlocking new data modalities and algorithmic techniques over a much larger range of diverse inputs. However, these methodological and technological improvements coupled with the ever increasing space of varying embedding inputs and potential downstream tasks have made it difficult to determine the optimal embedding strategy for a problem of interest. Thus far, current embedding benchmarking studies carve up this large space by examining only a single input data type or narrowly defined task, such as drug response [14] or biological function prediction [15], leaving a critical gap in understanding how embedding methods perform across a broader range of gene-associated applications. While embedding methods for protein-centric tasks have been more thoroughly characterized [7], there is still a need for comprehensive guidelines to inform the use of gene embeddings for common functional prediction tasks.

Here, we evaluate 38 state-of-the-art gene embedding methods across 3 categories of benchmarking tasks designed to assess each method’s performance in predicting: (1) individual gene level attributes and disease genes, (2) paired gene interactions and pathway edges, and (3) relationships between functionally meaningful collections of genes. Critically, our gene embedding benchmarking suite assesses performance across a variety of functional downstream tasks, while accounting for differences in gene coverage and data leakage. We find that the underlying input data type is the most critical factor influencing performance, with biomedical literature-based embeddings, particularly GenePT-Model3 [16], demonstrating strong generalizability across all tasks. Other embeddings perform well in specific scenarios, such as amino acid–based embeddings for predicting regulatory relationships, gene expression-based embeddings for disease-related tasks, and embeddings explicitly encoding gene relationships, such as protein-protein interaction networks, typically perform well on functional and genetic interaction prediction tasks.

## Results

### Benchmarking suite for assessing gene embeddings on functional classification tasks

We reviewed and categorized 38 publicly available gene embedding methods based on their input data types, algorithms, and embedding dimensionalities (Table 1). These embeddings span five broad data types, including amino acid sequences, biomedical literature, gene expression (single-cell and bulk RNA-seq), mutation profiles, and protein-protein interactions (PPI). We further categorized the embeddings by their construction algorithm into six categories: non-machine learning methods (non-ML), matrix factorization, multi-layer perceptrons (MLPs), recurrent neural networks (RNNs), skip-gram/CBOW, and transformers. Non-machine learning methods are a catch-all category for classic vector representations of genes with minimal further processing, such as protein sequence information from BLAST, PFAM, HMMER, or the raw PPI adjacency matrix. These serve as an important complement to ML-based embeddings and have historically been used in function classification tasks [36].

**Table 1.** Embedding Methods Tested. A summary of the embedding methods evaluated in this study, including their training input data types, underlying algorithm class, and dimensionalities. For each method, the number of genes (with mapped Entrez ids) and number of embedding dimensions are shown in parentheses.

| Embedding Name | Training Input Data | Algorithm | Dimensions |
| --- | --- | --- | --- |
| AAC [7, 11] | Amino acid sequence | Non-ML | (18,910, 20) |
| APAAC [7, 12] | Amino acid sequence | Non-ML | (18,908, 80) |
| BLAST [7, 8] | Amino acid sequence | Non-ML | (18,910, 20,421) |
| HMMER [7, 9] | Amino acid sequence | Non-ML | (18,910, 20,421) |
| KSEP [7, 13] | Amino acid sequence | Non-ML | (18,835, 400) |
| PFAM [7, 9] | Amino acid sequence | Non-ML | (18,910, 6,227) |
| CPC-PROT [7, 17] | Amino acid sequence | RNN | (18,493, 512) |
| SEQVEC [7, 18] | Amino acid sequence | RNN | (18,910, 1,024) |
| UNIREP [7, 19] | Amino acid sequence | RNN | (18,910, 5,700) |
| LEARNED-VEC [7, 20] | Amino acid sequence | skip-gram/CBOW | (18,910, 64) |
| PROTVEC [7, 21] | Amino acid sequence | skip-gram/CBOW | (18,910, 100) |
| ALBERT [7, 22] | Amino acid sequence | Transformer | (18,493, 4,096) |
| BERT-BFD [7, 22] | Amino acid sequence | Transformer | (18,493, 1,024) |
| BERT-PFAM [7, 23] | Amino acid sequence | Transformer | (18,494, 768) |
| ESM2 [24] | Amino acid sequence | Transformer | (18,902, 1,280) |
| ESMB1 [7, 25] | Amino acid sequence | Transformer | (18,493, 1,280) |
| ProtT5 (T5) [7, 22] | Amino acid sequence | Transformer | (18,493, 1,024) |
| XLNET [7, 22] | Amino acid sequence | Transformer | (18,493, 1,024) |
| BIOCONCEPTVEC-GLOVE [26] | Biomedical literature | Matrix factorization | (20,917, 100) |
| BIOCONCEPTVEC-CBOW [26] | Biomedical literature | skip-gram/CBOW | (20,917, 100) |
| BIOCONCEPTVEC-FASTTEXT [26] | Biomedical literature | skip-gram/CBOW | (20,917, 100) |
| BIOCONCEPTVEC-SKIP-GRAM [26] | Biomedical literature | skip-gram/CBOW | (20,917, 100) |
| GenePT-ADA [16] | Biomedical literature | Transformer | (33,837, 1,536) |
| GenePT-Model3 [16] | Biomedical literature | Transformer | (33,837, 3,072) |
| FROGS-ARCHS4 [27] | Gene expression (bulk) | MLP | (22,970, 256) |
| TCGA-EMBEDDING [7, 28] | Gene expression (bulk) | MLP | (18,432, 50) |
| Gene2Vec [7, 29] | Gene expression (bulk) | skip-gram/CBOW | (17,798, 200) |
| GF-12L30M [30] | Gene expression (single cell) | Transformer | (21,228, 512) |
| GF-12L95M [30] | Gene expression (single cell) | Transformer | (19,523, 512) |
| GF-12L95MCANCER [30] | Gene expression (single cell) | Transformer | (19,523, 512) |
| GF-20L95M [30] | Gene expression (single cell) | Transformer | (19,523, 896) |
| GF-6L30M [30] | Gene expression (single cell) | Transformer | (21,228, 256) |
| scGPT-human [31] | Gene expression (single cell) | Transformer | (38,714, 512) |
| scGPT-pancancer [31] | Gene expression (single cell) | Transformer | (38,714, 512) |
| Mut2vec [7, 32] | Mutation profile, Literature, PPI | skip-gram/CBOW | (17,500, 300) |
| Mashup [33] | PPI | Matrix factorization | (21,503, 800) |
| PPI-raw [34] | PPI | Non-ML | (20,363, 20,363) |
| Node2Vec [35] | PPI | skip-gram/CBOW | (20,362, 128) |

To facilitate the most common use cases, we used gene embeddings as published or extracted them with minimal preprocessing. Given the heterogeneity of input data and embedding methodologies involved, both the gene coverage and the dimensionalities of the embeddings vary highly. Total gene coverage ranges from 17,500 genes at the low end (Mut2Vec [32]) up to 38,714 genes (scGPT [31]; Table 1). To account for differences in gene coverage, we developed two variants for each benchmarking task, one with the common set of 11,355 genes shared across all embeddings tested and one with the complete embeddings referred to as “full” in our tests. Embedding dimensionality ranges from compact vectors as low as 20 dimensions (AAC [7, 11]) to detailed representations up to 20,421 dimensions (BLAST [7, 8] and HMMER [9]; Table 1). In all our benchmarking tests, we use the entire set of embedding dimensions, with no reduction.

Evaluating gene embeddings across a spectrum of biological tasks while isolating embedding quality from downstream model complexity is challenging. To address this, we designed a benchmarking suite with three complementary evaluation tracks that together probe different aspects of embedding representations: (1) prioritizing single gene attributes, (2) predicting paired interactions, and (3) comparing gene sets (Figure 1). For single and paired gene tasks, we use support vector machine (SVM) classifiers with a radial basis kernel to perform supervised learning across 5 core functional, disease, and genetic interaction prediction tasks. Using a classifier like SVM makes it easier to isolate the “quality” of an embedding from idiosyncrasies of more complex model architectures. Furthermore, SVMs are particularly well-suited to handling diverse embedding dimensions and modalities, minimizing downstream model bias and overfitting in high-dimensional, low-sample settings while keeping the hyperparameter tuning process simple and fast. To compare gene sets, we use our previously developed method, ANDES [37], which efficiently ranks gene sets based on their similarity within an embedding space. We use the set matching ability of ANDES to rank gene embeddings based on their performance in matching known functionally similar gene sets and paired disease-tissue gene sets.

**Figure 1.**
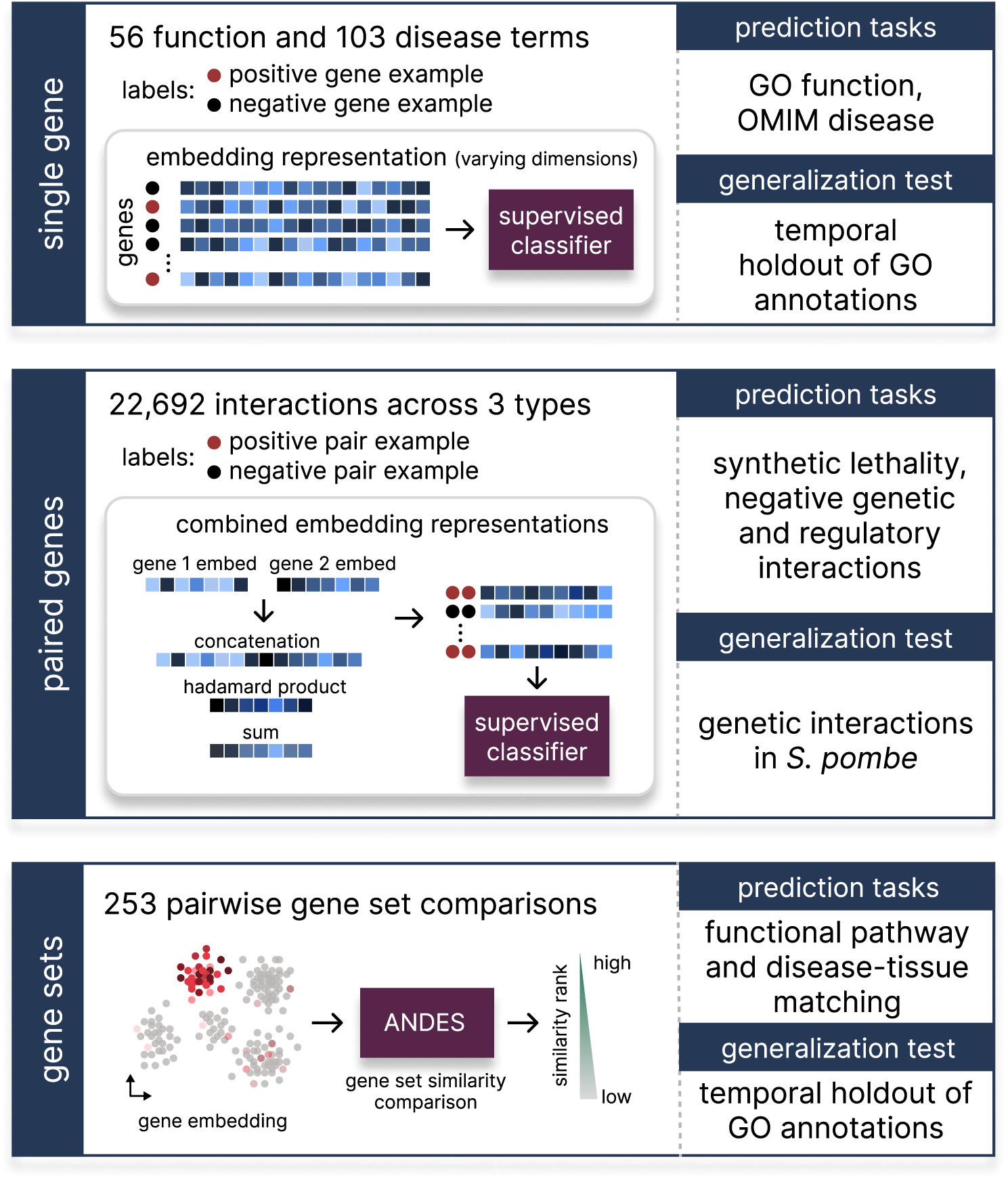
Overview of benchmarking tasks and classification setup. 38 embedding methods using different underlying data and construction methodologies were evaluated across three classes of common prediction tasks: single gene tasks determining gene function or identifying disease genes, paired gene tasks identifying synergistic or antagonistic effects involving 2 genes, and gene sets tasks where embeddings are used to measure similarity between functionally or mechanistically similar sets of genes.

While our benchmarking suite is structured to measure different gene relationships and accounts for the multi-functionality of genes, the diverse data inputs used in constructing gene embeddings still create possible data leakage, particularly in settings where tasks involve the prediction of shared gene pathways or well-known annotations. Data leakage is less of a problem for methods that train primarily on a single data type, such as amino acid sequence and gene expression-based models, or even protein-protein interaction networks, but is potentially concerning in embeddings derived from biomedical literature and multi-input models that may use functional annotations. To mitigate this, we added a dedicated temporal holdout task for the single gene and gene set evaluations, particularly gene ontology (GO) annotations that were released after the latest embedding method we evaluate was trained (March 2024). Furthermore, for predicting pairwise gene relationships, we created an evaluation set of genetic interactions for a less-studied organism *Saccharomyces pombe (S. pombe)* which, to the best of our knowledge, was not used as input for any embedding method. Methods that perform well on these tasks are likely to be more generalizable.

### Gene-level attribute benchmarks

To evaluate how well each embedding captures *individual gene-level attributes*, we performed two primary classification tasks: gene function prediction using GO [38] terms and disease-gene prediction using OMIM [39] annotations. For each GO or OMIM term, we trained a separate SVM classifier for every embedding, using the embedding vectors as input features to predict held-out genes sharing that annotation (Figure 1). To focus on distinct, nonredundant biological categories and reduce overlapping or hierarchical annotations, we curated “slim” sets of GO Biological Process and Disease Ontology [40] terms (see Methods). Additionally, to prevent circularity and minimize data leakage, we included only GO annotations supported by experimental evidence codes and excluded any GO or OMIM term associated with fewer than 20 genes to ensure adequate sample size for training and testing. After these filters, the benchmark comprised 56 GO terms and 103 OMIM disease terms, with an additional 19 GO terms reserved as a temporal holdout to approximate generalizability.

Across both gene function and disease-gene prediction tasks, literature-based embedding methods consistently outperformed those trained on other input data types (Figure 2). GenePT-Model3 [16] was the top performer in both settings (mean AUROC GO: 0.926; OMIM: 0.877), followed by the BioConceptVec [26] family of models. Notably, BioConceptVec-FastText achieved comparable performance (mean AUROC GO: 0.909; OMIM: 0.871) to GenePT-Model3 while using far fewer dimensions (Table 1), making BioConceptVec much more computationally efficient, approximately 22.3 times faster in our tests (Table S1). The BioConceptVec-GloVe variant performed notably worse relative to other literature embeddings (rank 12; mean AUROC GO: 0.799; OMIM: 0.721).

**Figure 2.**
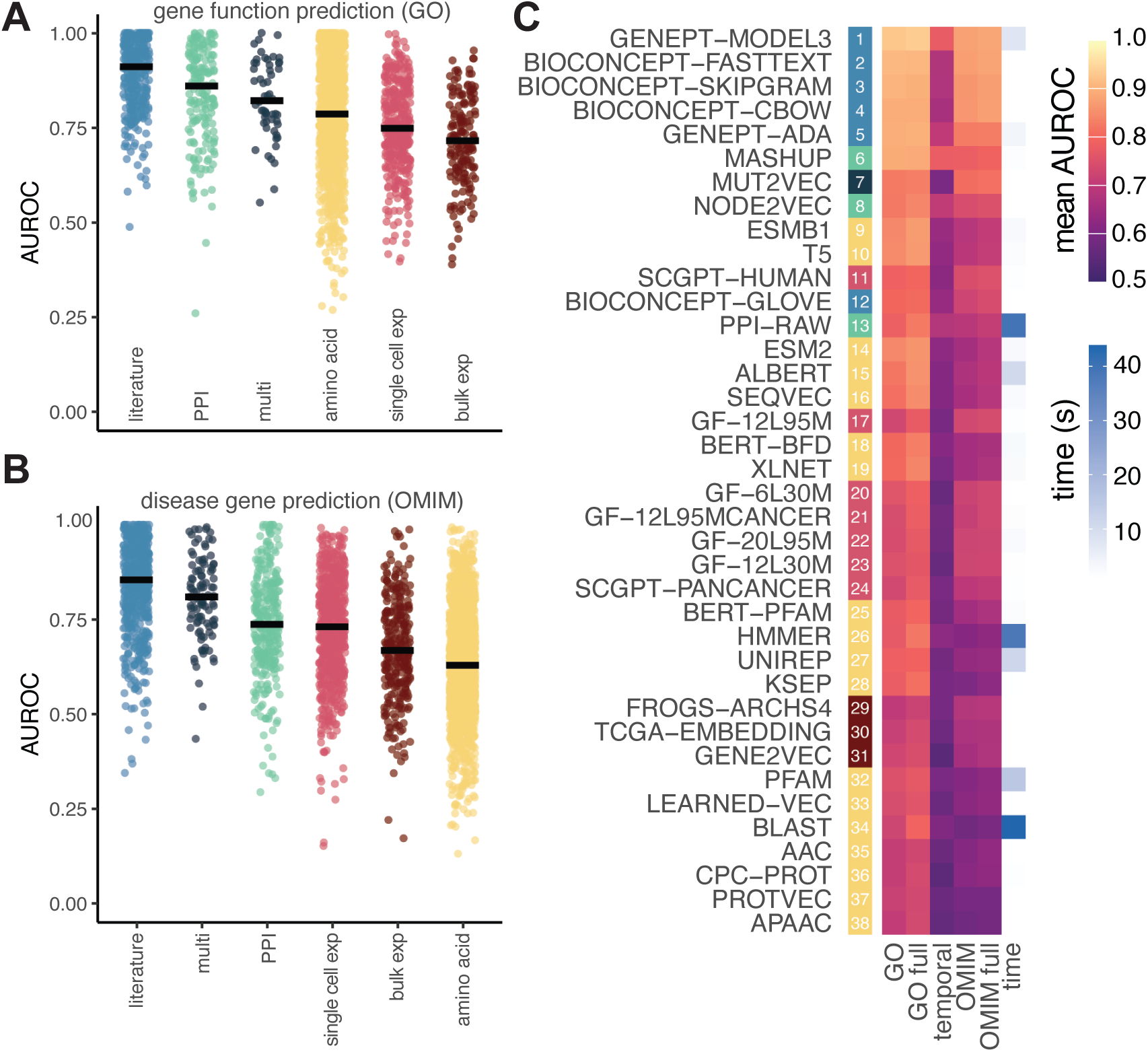
Benchmarking results for single gene tasks. Embedding methods were evaluated for their performance as features in predicting two classes of disease attributes: (A) gene function using GO and (B) disease gene identification using OMIM. In both cases embeddings are categorized and sorted by the median performance of their primary input data type with each dot representing one AUROC score for a single embedding type and GO term (A) or disease (B) pair. (C) Summarizes findings by ranking embedding methods according to the mean AUROC across tasks. Columns denoted as “full” use the entire set of genes covered in the released embeddings. The results for the GO temporal holdout is summarized in the temporal column. The time column highlights the differences in average training times for a single SVM classifier using the different embeddings.

PPI-based embeddings occupied the next performance tier, with Mashup [33] and Node2Vec [35] outperforming sequence– and expression-based models (mean AUROC Mashup GO: 0.891, OMIM: 0.782; Node2Vec GO: 0.824, OMIM: 0.742). Mashup performed best among network-derived methods and was also the most stable under temporal holdout (see below). We also evaluated Mut2Vec [32], a multimodal embedding trained jointly on PPI, somatic mutation profiles, and co-occurrence in biomedical literature. Mut2Vec was on par with PPI-based embeddings on GO but surpassed them on disease-gene prediction (mean AUROC GO: 0.824; OMIM: 0.805), suggesting that incorporating cancer-related mutation data and textual context provides a modest, biologically intuitive advantage for identifying disease-associated genes.

Amino-acid and expression-based embeddings generally performed below the literature– and network-derived models (Figure 2A,B), though several sequence models approached mid-tier performance. In general, sequence-derived embeddings tended to perform better for gene-function prediction than for disease-gene prediction, consistent with their emphasis on molecular and structural features. Among them, ESMB1 [7, 25] and ProtT5 [7, 22] were competitive with the PPI-based embeddings (Figure 2C), ranking just below Node2Vec and above BioConceptVec-GloVe (mean AUROC, ESMB1: GO: 0.841, OMIM: 0.685; ProtT5: GO: 0.845, OMIM: 0.691).

Expression-based models showed the inverse trend, performing significantly better than the sequence-based models for disease-gene prediction (paired Wilcoxon signed-rank test, single cell RNA-seq (scRNAseq) vs amino acid: *p* = 1.34*e*^−9^; bulk vs amino acid: *p* = 3.72*e*^−3^). Although still trailing the top tier of performers, single-cell embeddings approached PPI-based performance in this task (mean AUROC OMIM PPI: 0.74; scRNAseq: 0.72, paired Wilcoxon signed-rank test, *p* = 0.73) and exceeded bulk expression embeddings (mean AUROC OMIM: 0.72 vs 0.67; paired Wilcoxon signed-rank test, *p* = 1.83*e*^−8^). scGPT-human [31] achieved the top overall performance among expression-based methods (mean AUROC GO: 0.787; OMIM: 0.744). In general, we see that while sequence and expression embeddings remain below the literature and PPI tiers, certain sequence models (ESMB1, ProtT5) and expression models (scGPT-human) are still competitive. Interestingly, we see the same general trend where structure-focused models favor gene function prediction, whereas context-aware representations better capture disease associations.

In the temporal holdout task (“temporal” column, Figure 2C), classifiers were trained on pre-2024 annotations and tested on genes annotated after March 2024. Because none of these updated gene annotations were included in the training data for any embedding method, this evaluation provides a leakage-free measure of generalization. As expected, performance declined in this more challenging setting (mean AUROC across all methods dropped from 0.79 to 0.62). Despite this drop, the relative ranking of models remained consistent, with PPI-based embeddings (Mashup, Node2Vec, PPI-Raw) showing the strongest generalization, performing better than literature-based models (mean AUROC GenePT-Model3: 0.774; Mashup: 0.777; Node2Vec 0.704; PPI-Raw 0.682). Performance varied across individual GO and OMIM terms, with some (e.g., “endodermal cell differentiation”, “congenital disorder of glycosylation”) consistently easier to predict, while others (e.g., “glucose homeostasis”, “spinal disease”, and “motor neuron disease”) being more difficult across all models (Figures S1 and S2).

### Paired gene interaction benchmarks

Next, we assessed the utility of gene embeddings in modeling *pairwise gene interactions*, a common use case for systematic hypothesis generation in genetic, drug perturbation, and other functional screens. Predictive models that accurately capture these relationships can help prioritize gene pairs for combinatorial perturbation, from dual knockdowns to co-inhibition strategies, depending on the desired experimental or therapeutic goal. We evaluated each embedding method as input features for classifiers across three distinct interaction types: negative genetic interactions (NG), synthetic lethal (SL), and transcription factor-target (TF) pairs (Figure 3). Since embeddings are at the individual gene level, we constructed pairwise representations using 3 different operations: concatenation (cat), element-wise sum (sum), and Hadamard product (prod). We then trained separate classifiers for each interaction and operation type. Unlike the single gene annotation models which train a separate classifier per GO or OMIM term, each pairwise model predicts interactions across all gene pairs using nested cross-validation (Figure 1).

**Figure 3.**
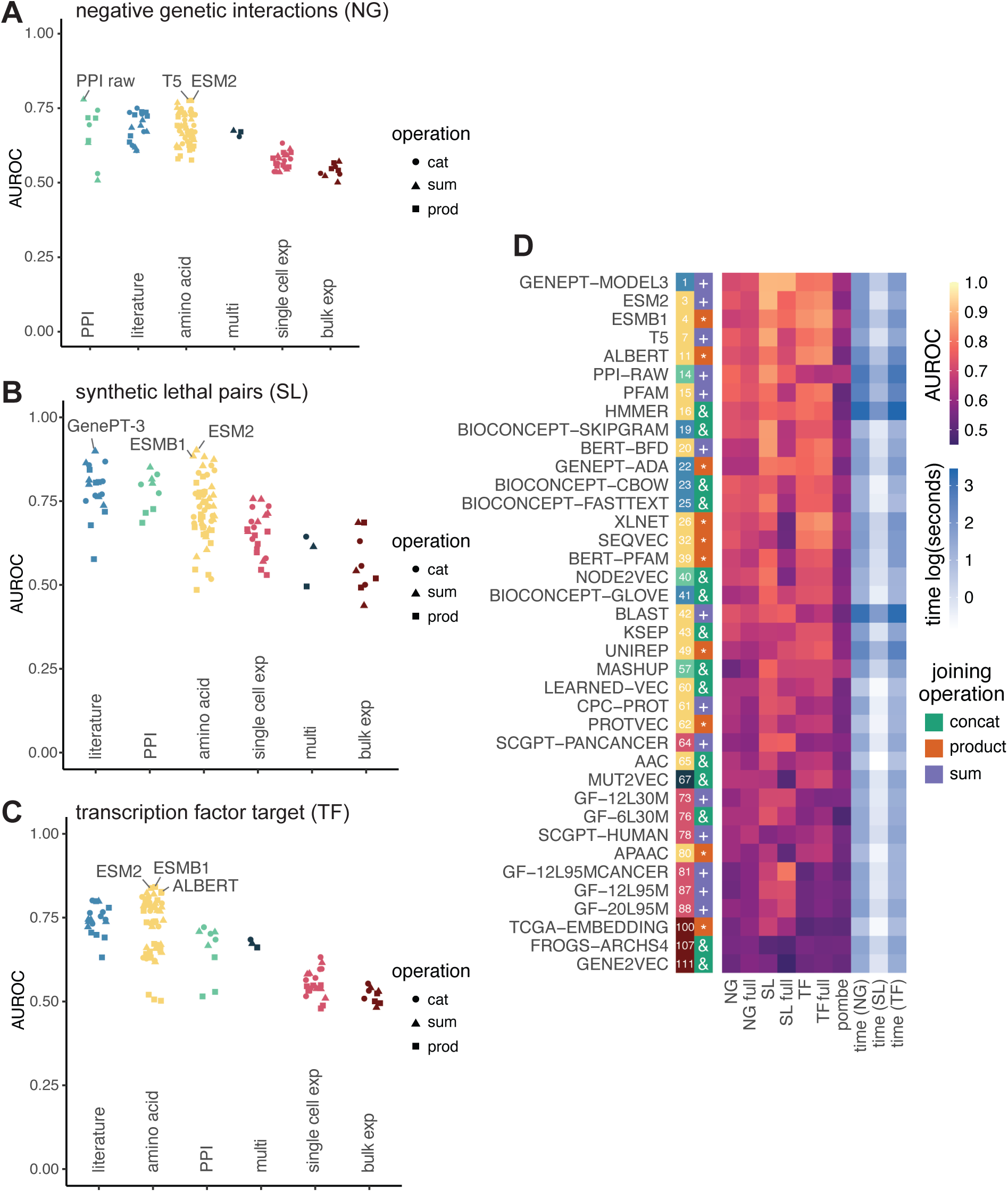
Benchmarking results for paired gene tasks. Embedding methods were evaluated based on their effectiveness as features in three main prediction tasks: (A) negative genetic interactions, (B) synthetic lethal pairs (B), and (C) transcription factor–target interactions. All tasks involve gene pairs, and we assessed three strategies for combining embeddings from each gene in a pair: concatenation (circles, labeled “cat”), element-wise sum (triangles), and Hadamard product (squares, labeled “prod”). Methods are grouped by the input data type used to generate embeddings. For each task in (A-C) the top 3 performing methods are labeled. Results are summarized in (D): the first column ranks methods by overall performance across the three within-species tasks (colored by input data type), with the best-performing pairwise combination shown in the second column. The next three columns display AUROC values for each task, followed by generalization performance to S. pombe. Columns labeled “full” refer to models trained on the full set of genes covered by each embedding. The final three columns report training times for each task (log-transformed seconds), where darker shading indicates longer training times.

SL prediction proved to be the most tractable of the three tasks (mean AUROC across all methods and operations SL: 0.71, NG: 0.65, TF: 0.67; Figure 3), likely reflecting its well-defined phenotypic endpoint of cell death. NG represent a broader set of genetic interactions ranging from moderate to severe, and TF regulatory interactions can vary depending on cellular context, likely making these two tasks more challenging. Across tasks, the same general set of high-performing embeddings (GenePT-Model3, ESM2, ESMB1, T5, ALBERT, PPI-raw) reappeared, though their relative ranking varied by interaction type (Figure 3A-C). Ranking overall performance across all tasks and the *S. pombe* generalization test (a cross-species benchmark using a less extensively studied yeast to assess generalizability beyond well-characterized systems), the literature-based GenePT-Model3 embedding narrowly outperformed the amino acid-based ESM2 [24] and ESMB1 [25] (mean AUROC across tasks for top-performing operation, GenePT-Model3: 0.777, ESM2: 0.765, ESMB1: 0.762; Figure 3D). Generally, these three methods had comparable performance, with GenePT-Model3’s slight edge likely stemming from its increased gene coverage (number of genes, GenePT: 33,837, ESM2: 18,902, ESMB1: 18,493) in the full embeddings for the NG, SL, and TF tasks.

Compared with the single-gene benchmarks, sequence-based embeddings performed substantially better in these pairwise interaction tasks, often matching or exceeding the literature-derived models. We reason that this is possibly because protein-level representations directly encode biophysical compatibilities that underlie genetic and regulatory interactions. In contrast, PPI-based embeddings, which were among the top performers for NG and SL, declined on the TF task. This likely reflects the fact that TF-target relationships in this benchmark capture protein-DNA regulatory interactions, which may be informed by, but are distinct from the protein-protein associations encoded in the PPI-based embeddings.

Several non-ML based embeddings (PPI-raw [34], PFAM [9], and HMMER [9]) also performed unexpectedly well (mean AUROC across tasks for top operation, PPI-raw: 0.724, PFAM: 0.723, HMMER: 0.721). PPI-raw ranked among the top three methods for NG classification (AUROC=0.779; Figure 3A) and achieved the highest performance in the *S. pombe* generalization task (AUROC=0.653; Figure 3D). Mashup also performed well in this cross-species evaluation (AUROC=0.646), ranking second behind PPI-raw, highlighting the robustness of network-based representations that we also observed in the single-gene benchmarks.

The choice of operation for combining embeddings into pairwise representations significantly influenced performance across tasks (ANOVA, F-value= 16.114, *p* = 3.0*e*^−7^), albeit less impactful than the embedding input data type (ANOVA, F-value= 89.429, *p <* 2*e*^−16^) or embedding construction algorithm (ANOVA, F-value= 23.570, *p <* 2*e*^−16^). Generally, the product operation yielded lower performance than concatenation or summation (ANOVA, prod vs cat: *p.adj* = 0.009, prod vs sum: *p.adj* = 0.007), with this effect most pronounced among non-ML-based embeddings (ANOVA, prod vs cat: *p.adj* = 0.001, prod vs sum: *p.adj* = 0.001; Figure S3), though some embedding methods (e.g., ESMB1, ALBERT, GenePT-ADA) showed the opposite trend. Given these results, we recommend the element-wise summation operation as a general default, as it provides similar performance to concatenation, but with substantially shorter training and prediction times (Table S1).

### Gene set comparison benchmarks

Building upon our evaluations of individual genes and gene pairs, we next extended our analysis to *gene sets*, which describe collective properties of functionally related genes. Whereas the single gene benchmarks evaluated how well embeddings capture individual membership with known functional or disease categories (Figure 2) and the pairwise benchmarks tested their ability to model gene-gene relationships (Figure 3), this analysis examined whether the embedding spaces themselves effectively encode higher-order relationships between groups of genes, such as shared pathway membership or cross-disease and tissue associations. To facilitate this, we used our previously developed tool, ANDES [37], which computes reciprocal best-match similarities between genes in two sets and aggregates them into a single score. This approach preserves the diversity of relationships within gene sets and corrects for biases due to set size, enabling comparisons even between entirely non-overlapping sets. We evaluated three types of gene set relationships: KEGG-GO pathway alignment, disease-tissue associations, and a temporal GO holdout to evaluate generalizability.

For the KEGG-GO matching task, we evaluated how effectively each embedding recovered 54 curated KEGG-GO term pairs representing the same biological processes [37] (Figure 4A). To avoid artificial inflation of similarity scores, genes shared between each pair were removed from the GO terms, ensuring that KEGG and GO sets were non-overlapping. All embedding classes performed reasonably well at this task, with literature– and PPI-based embeddings achieving the highest average performance (mean normalized rank (nrank), literature: 0.927, PPI: 0.883; Figure 4A). Among individual methods, GenePT-Model3 was again the top performer (mean nrank: 0.951), closely follwed by GenePT-ADA (mean nrank: 0.941) and other literature models, BioConceptVec CBOW (mean nrank: 0.936) and skip-gram (mean nrank: 0.923). The PPI-based Node2Vec embedding completed the top 5 with performance comparable to the leading literature-based embeddings (mean nrank: 0.921).

**Figure 4.**
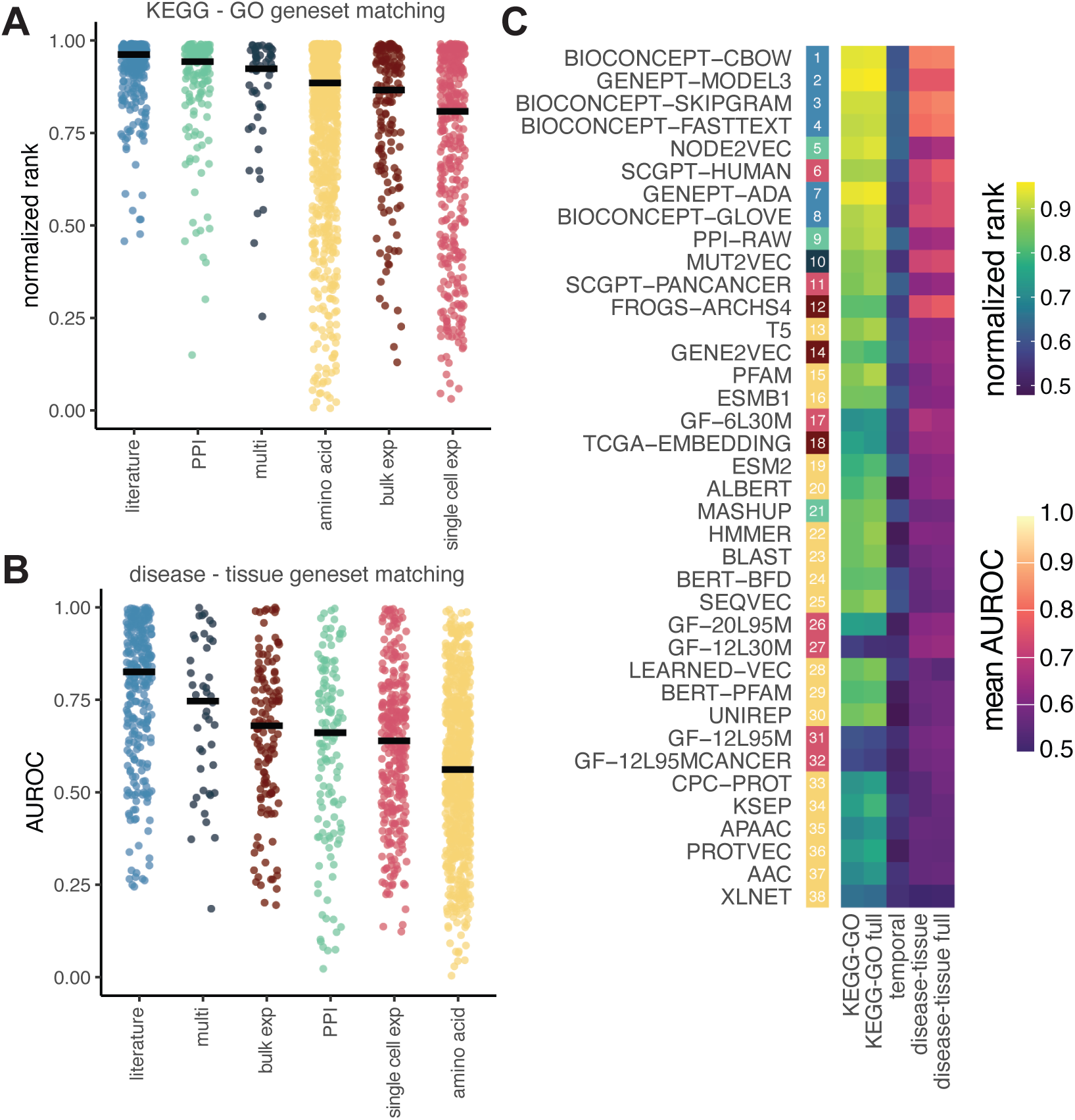
Benchmarking results for gene sets. Embedding methods were evaluated for their performance in gene set matching tasks using ANDES. In the first task (A), KEGG gene sets were matched to corresponding GO pathways, with results displayed as normalized ranks (computed as 1 – rank / all possible matches). In the second task (B), disease-associated gene sets were matched to relevant tissue gene sets, and performance was assessed using AUROC to quantify recovery of correct disease–tissue associations. Results are summarized in (C) where methods are ranked by their overall performance across the KEGG-GO and disease-tissue matching tasks (first column). As in previous figures, columns labeled “full” indicate models trained on the entire set of genes represented in each embedding. The “temporal” column highlights the potential generalization capacity of each embeddings by testing its ability to match the same GO gene sets (without overlapping genes) across a temporal holdout.

For the disease-tissue gene set matching task, we assembled a set of 200 disease-tissue term pairs based on manual curation of OMIM descriptions linking disease phenotypes to their tissue manifestations, encompassing 44 tissue and 79 disease gene sets. This task assesses how well embeddings capture latent biological relationships connecting disease-associated and tissue-specific genes. Literature-based embeddings again achieved the highest average performance, whereas PPI-based methods dropped substantially, performing below embeddings built from bulk expression data (mean AUROC, literature: 0.773, multi: 0.716, bulk exp: 0.664, PPI: 0.612; Figure 4B). Among literature models, BioConceptVec CBOW (mean AUROC 0.827) and fastText (mean AUROC 0.802) surpassed GenePT-Model3 (mean AUROC 0.760). The bulk-expression embedding FRoGS-ARCHS4 [27] was the top non-literature performer, with performance approaching GenePT-Model3 (mean AUROC 0.741). Interestingly, bulk-expression embeddings performed substantially better in this disease-tissue task than in KEGG-GO matching or other benchmarks, perhaps reflecting their ability to capture tissue-level expression context. On the other hand, the decline in performance for PPI-based embeddings may be highlighting a potential limitation in modeling these types of context-specific relationships.

As with all of our benchmarking tasks, we aimed to evaluate generalizability by including test cases unlikely to be included as training for the original embedding models. Here, we again designed a temporal holdout of GO terms, using terms with annotation updates before and after March 2024 where a minimum of 5 genes were added. For each such term, we applied the ANDES gene set matching test between its pre– and post-update versions, reasoning that they should remain functionally similar across updates. Although matching performance varied widely across terms (Figure S4), PPI-based embeddings again emerged on top (mean nrank, PPI-raw: 0.634, Node2Vec: 0.634; Figure 4C). The highest performing embeddings for the KEGG-GO and disease-tissue matching task also performed well in this temporal task, with BioConceptVec skip-gram (mean nrank, 0.632) and fast-text mean nrank, 0.629) having comparable performance to the PPI embeddings.

Overall, the top ranked methods performed similarly across the three gene set tasks, with a few exceptions (Figure 4C). Literature-based embeddings remained consistently strong across all benchmarks, reflecting their broad robustness across biological contexts. PPI-based methods, especially Node2Vec and PPI-raw, showed standout performance for the pathway matching tasks. The expression-based methods performed well on the disease-tissue association task (e.g., scGPT-human, FRoGS-ARCHS4), highlighting the added value of tissue-level context. The amino acid models were generally less effective for gene set matching, although ProtT5 (T5), PFAM, and ESMB1 were the strongest within this category.

### Determining influential embedding attributes

To better understand what drives differences in embedding performance across tasks, we examined how embeddings relate to each other in terms of their gene coverage and embedding similarity. While most embeddings covered largely overlapping genes (average Jaccard index = 0.796; Figure S5), recent transformer-based models such as scGPT and GenePT represented substantially more genes overall (38,714 and 33,837, respectively, versus ∼17,500-23,000 genes for other methods) and showed lower overlap with other embeddings (mean Jaccard index, scGPT models = 0.517; GenePT models = 0.573), but had high overlap with each other in gene coverage (Jaccard index = 0.865). Among the remaining methods, FRoGS-ARCHS4 and Mashup had somewhat lower overlap with other embeddings (mean Jaccard index, FRoGS = 0.599; Mashup = 0.620), although this difference in coverage did not seem to confer any additional advantage when using their non-intersecting (“full”) embeddings (Figures 2 to 4).

Beyond gene coverage, we assessed embedding similarity using canonical correlation analysis (CCA) across all shared genes (Figure 5A). Because CCA does not require identical vector dimensionality, it enabled direct comparison among the diverse embedding methods. Embeddings clustered primarily by their input data type, but with some notable deviations often corresponding to algorithmic differences. For example, several non-ML sequence-based models derived from alignment or motif methods (APAAC, BLAST, HMMER, PFAM) were largely uncorrelated with the learned amino acid embeddings. The PPI-based models (Mashup, Node2Vec, PPI-RAW) were also distinct from one another and other data types, with Node2Vec occupying an intermediate position within a broader cluster of literature-based methods, while Mashup and PPI-RAW were more isolated from all other models. Expression-based embeddings from bulk and single-cell data clustered moderately together, reflecting some of the shared transcriptome-based signal despite differences in underlying data compendia. As expected, variants within the same model family (e.g., GenePT, BioConceptVec, scGPT, GeneFormer) clustered more so with each other. This organization highlights the strong influence of data type and model family on learned embedding structure.

**Figure 5.**
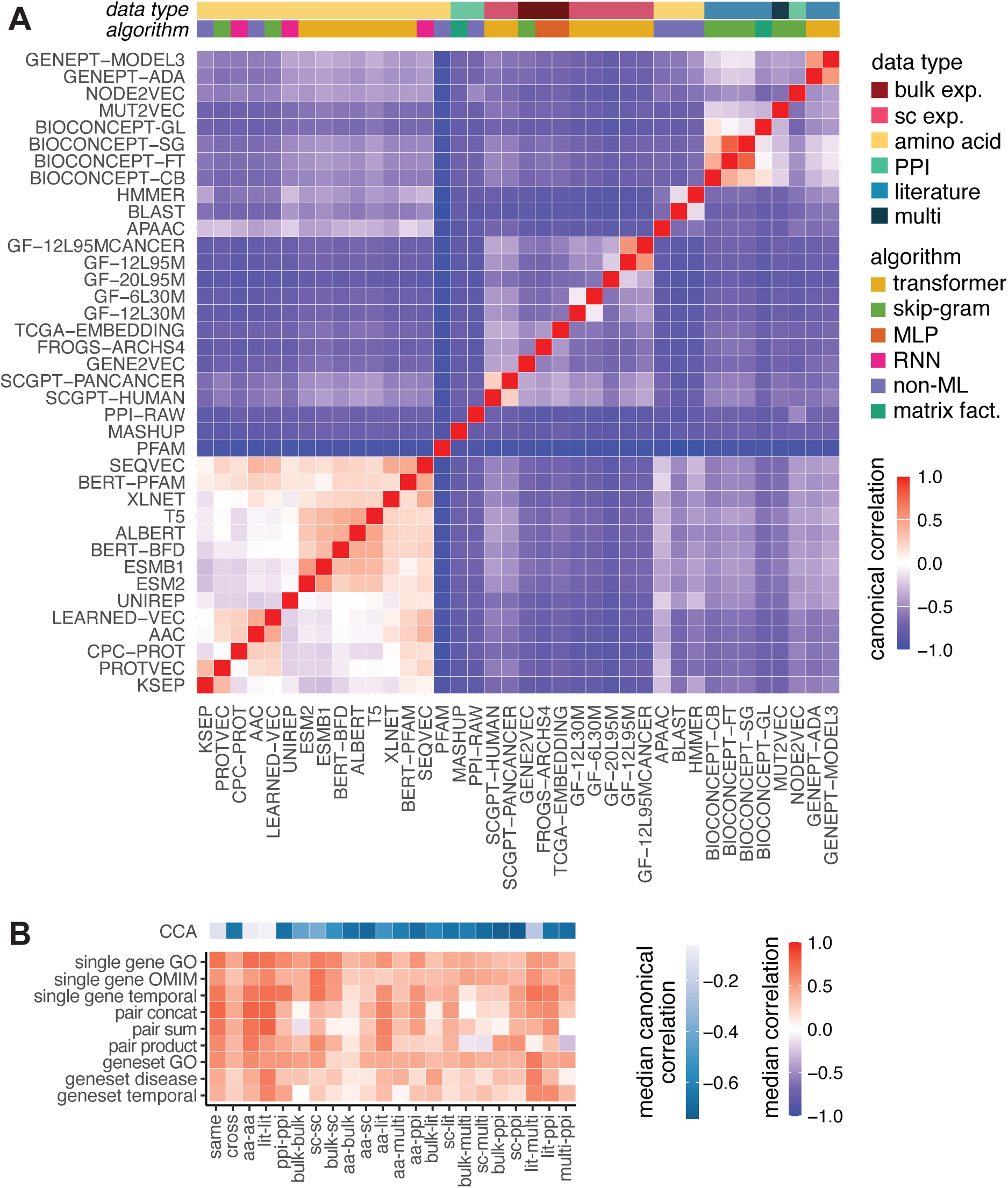
Similarity of embeddings and performance summary. (A) Clustered heatmap showing canonical correlation analysis (CCA) values between embedding methods, capturing the similarity structure of learned representations. Colored bars annotate each method by its input data type and embedding algorithm. (B) Heatmap illustrating how embedding similarity relates to performance across the single-gene and gene set tasks. The top bar summarizes CCA values by computing the median similarity within and between input data type groups. Pairwise Spearman correlation coefficients were calculated on task performance and were aggregated by embedding input data type pairs using the median to highlight general trends. When methods have the same input data type (e.g., literature-literature, PPI-PPI) we summarize them as “same” and when method pairs are different types (e.g., literature-PPI, amino acid-PPI) we provide an overview as “cross”.

Unsurprisingly, data type was the most influential factor influencing performance across all benchmarks (ANOVA across all tasks, *p <* 0.01; Table S2). The algorithm used to construct each embedding mattered most for the single and paired gene tasks, accounting for up to 32% of the total variance depending on the pairwise joining operation. Meanwhile, embedding dimensionality had a negligible impact on these same tasks (0.3%-6% of the variance), but had a significant impact for the gene set tasks (ANOVA p, KEGG-GO: 0.010; disease-tissue: 0.002; Table S2), explaining 19.79% of the total variance on the KEGG-GO matching and 22.39% on the disease-tissue matching task. These findings highlight that while input modality and algorithm are the most important for overall performance, embedding size begins playing a larger role in higher-order, multi-gene evaluations.

To explore the extent to which embedding similarity is reflected in performance similarity, we correlated the task-level prediction metrics between every pair of models across benchmarking tasks (Figure 5B). As expected, median correlations were high among methods derived from the same input data type, while cross-type correlations were lower. Interestingly, expression-based embeddings, particularly those derived from bulk data, showed weaker within-type correlations, suggesting greater heterogeneity among these models. Correlations were also generally lower for the multi-gene tasks (pair and gene set benchmarks), consistent with reduced performance concordance as the predictive context becomes more complex.

## Discussion

In this study, we undertook a comprehensive benchmarking effort to evaluate gene embeddings across a wide range of gene-centered prediction tasks. One of the most notable findings is the strong performance of text-based embeddings and the overall weak performance of expression-based models across all tasks. GenePT-Model3 was among the top performers in nearly all evaluations except the disease-tissue gene set matching task, which is the least well-characterized. The impressive performance of text-based embeddings raises concerns about potential data leakage, since these embeddings are trained on curated documents or vast corpora of biomedical literature that may implicitly contain knowledge overlapping with our evaluation benchmarks. While it is challenging to completely separate the collective knowledge of biomedical research from our gold standards, we attempted to mitigate it with carefully designed temporal and organism-specific holdouts. In these benchmarks, we did not observe any major performance drops that would indicate substantial leakage. Interestingly, PPI-based methods performed especially well in these evaluations. We speculate that their explicit encoding of gene-gene relationships may enhance their generalizability to unseen data.

Our analyses also further clarified the relative value of newer representation learning approaches versus their classical or non-machine learning counterparts, particularly for amino acid sequence embeddings, where methodological diversity is high. For this class of data, machine learning derived embeddings were generally more compact and computationally efficient (Table S1) while achieving higher performance. Transformer models such as ProtT5, ESMB1, and ESM2 consistently outperformed other sequence embeddings across tasks. Among the non-machine learning approaches, PFAM embeddings had the strongest overall performance, especially for predicting genetic interactions and modeling gene set relationships. Given its performance and availability, PFAM representations can serve as a useful baseline when evaluating future amino-acid sequence embeddings. Similarly, the PPI-raw embedding, which is simply the adjacency matrix of a consensus protein-protein interaction network, also performed competitively, often surpassing many sequence– and expression-based models despite its simplicity. It thus serves as not only a valuable baseline for network-derived embeddings, but also as an interpretable reference approach for other embedding efforts that aiming to model gene functional similarity.

One potential limitation of our study is the use of SVM models as the sole predictive model for evaluations. We adopted this design to isolate the information content of each embedding rather than optimize for classification accuracy. Nonetheless, it is possible that additional hyperparameter tuning, alternative kernels, or other classification algorithms could be better suited for particular embedding architectures or input data types. However, given that most of our test cases had a small to medium set of balanced labels, we believe that SVMs are a reasonable choice for cross-method comparison. In addition, the embeddings included here were trained independently using different datasets, corpora versions, and preprocessing pipelines, so performance differences may partly reflect disparities in data quality or training scope rather than intrinsic differences between data type and algorithm.

Beyond specific performance trends, our benchmarking framework provides a natural hierarchy in how biological information can be represented and evaluated. The three benchmarking categories, from single gene to gene pairs and gene sets, reflect increasing levels of biological abstraction, from molecular properties to interaction networks and collective pathway behavior. Viewing evaluation tasks through this lens clarifies which embedding modalities are best suited to different biological scales and underscores the importance of testing new models across this hierarchy.

In conclusion, our benchmarking study highlights the general versatility and robustness of literature-based embeddings, which achieved the most consistent high performance across the gamut of prediction tasks. Network-derived embeddings were competitive in several settings, especially with regards to gene function, but showed weaker results in others, such as transcription factor-target and disease-tissue prediction. Together, these insights provide a guide post for selecting and applying gene embeddings tailored to specific downstream analyses. An emergent theme across our evaluations is that embedding performance does not scale simply with data size or model complexity. Models built from structured, knowledge-rich resources, such as NCBI gene descriptions, curated interaction networks, or PFAM domain profiles, often performed better than those trained on much larger, unstructured corpora. For instance, GenePT-Model3, which combines a transformer architecture with NCBI gene descriptions, outperformed BioConceptVec models trained on a large corpus of biomedical literature, reflecting the joint influence of model design and input quality. Non-ML embeddings such as PPI-raw and PFAM achieved competitive performance, at times surpassing their counterparts built from the same data type, underscoring the value of using direct, signal-dense representations. This pattern may perhaps help explain the comparatively weak performance of expression-based embeddings, where sample diversity and signal heterogeneity make it challenging to derive stably meaningful gene-level representations. Looking ahead, we anticipate that future gene embeddings crafted through integrating structured biological knowledge with large-scale data, potentially through human-guided or hybrid learning frameworks, will yield more interpretable and biologically grounded representations for biomedical discovery.

## Methods

### Gene embedding acquisition and standardization

We performed a literature search for gene embedding methods and selected those that provided gene embeddings as part of their publication. After downloading, each embedding was processed through a standard pipeline, where all gene identifiers were converted to Entrez IDs using the mygene python library [41]. For genes with multiple identifiers corresponding to the same Entrez ID, the mean embedding was calculated to generate a single vector representation for the gene. Any genes that could not be mapped to Entrez IDs were excluded. Only embeddings with at least 15,000 Entrez genes were retained to ensure a sufficiently large and comparable intersection between embedding methods.

The embeddings AAC [11], ALBERT [22], APAAC [12], BERT-BFD [22], BERT-PFAM [42], BLAST [8], CPC-PROT [43], ESMB1 [25], GENE2VEC [29], HMMER [9], KSEP [13], LEARNED-VEC [20], MUT2VEC [32], PFAM [9], PROTVEC [21], SEQVEC [18], T5 [22], TCGA-EMBEDDING [28], UNIREP [19], and XLNET [22] were sourced from the Protein RepresentatiOn BEnchmark study [7]. The embeddings BioConceptVec [26], FRoGS [27], Geneformer [30], GenePT [16], Mashup [33], and scGPT [31] were retrieved from their respective publicly available repositories. In addition to these previously published embeddings, we generated 3 additional embeddings (Node2Vec, PPI-RAW, and ESM2). For Node2Vec and PPI-RAW, we started with a consensus human PPI network that combines experimentally-derived protein physical interactions from eight databases [34]. Node2Vec embeddings were generated using random walks on this network followed by skip-gram modeling, whereas PPI-RAW simply used the raw adjacency matrix of the consensus PPI as a direct embedding representation. We also generated gene-level embeddings from the pretrained ESM2 protein language model (esm2_t33_650M_UR50D) [24] using human protein sequences from Uniprot [44]. Following the recommended procedure from ESM2’s authors, we created gene-level embeddings via mean pooling aggregation of corresponding amino acid embeddings.

To enable comparability across methods and separate performance differences driven by gene coverage, all embeddings were filtered to a common set of genes by taking the intersection across all embeddings, resulting in a common set of 11,355 genes. This intersecting set is the default test case of our benchmarking suite, while the original embeddings with their full set of genes were evaluated separately and referred to as “full” in the benchmarks.

### Gene ontology gold standards

We used gene sets from the Gene Ontology (GO) [38] biological process (BP) namespace in both the single gene and gene set benchmarking tasks. To reduce potential circularity, only annotations supported by experimental evidence codes (EXP, IDA, IPI, IMP, IGI, IEP, HTP, HDA, HGI, and HEP) were included. Using the directed acyclic graph structure of the ontology, we also propagated gene annotations from child to parent terms through “is a” and “part of” relationships, such that final gene sets also contain genes annotated to their child term(s).

The gold standard for individual test cases was created by taking different subsets of the GO BP hierarchy. For the single gene analyses, we used the 2024-09-08 release of GO BP and retained GO terms with more than 20 annotated genes. To create a nonredundant but representative evaluation, we then filtered these terms to a “slim” set of 56 terms using the Alliance of Genome Resources (AGR) slim annotations by randomly sampling three GO terms per AGR term. To evaluate generalizability over time, we also constructed a temporal holdout for use in both the single gene and gene set benchmarks. GO annotations were dived into two parts: those released prior to 2024-03, and those released by 2025-03-16. We then identified terms that gained at least 5 genes between these versions, resulting in a final set of 19 GO terms for the temporal evaluation.

### OMIM disease gold standards

We used disease-gene annotations from Online Mendelian Inheritance in Man (OMIM) [39], downloaded in October 2023, for the single gene and gene set benchmarking tasks. OMIM identifiers were mapped to Disease Ontology (DOID) [40] terms, and child annotations were propagated to parent terms through the ontology hierarchy, following the same procedure used for GO. As with GO, we used the AGR slim to keep a final set of 103 OMIM disease terms.

### Paired gene interaction gold standards

Synthetic lethal (SL) and negative genetic (NG) interaction data were obtained from BioGRID [45] (4.4.240), and transcription factor-target relationships from the Transcription Factor Target Gene Database [46]. For the cross-species generalization benchmark, we used NG interactions from *Schizosaccharomyces pombe*, also obtained from BioGRID [45] (4.4.240). We mapped yeast genes to their human orthologs using orthology data downloaded from OrthoMCL [47] (v6.1).

### Gene-level benchmarks

For each GO or DOID term, we trained an independent Support Vector Machine (SVM) classifier using the scikit-learn package [48]. Positive examples were defined as genes annotated to the target gene set. To construct negative examples, we first excluded any genes belonging to a parent AGR slim for that set, then randomly sampled 10 times the number of positives across all non-parent slim terms, ensuring balanced representation from each slim term. We completely excluded 20% of the resultant gold standard as a holdout set. The remaining genes were used for three-fold cross-validation to tune the SVM C parameter within [0.1, 1, 10, 100, 1000] based on mean AUROC across folds. All models used the radial basis function kernel and balanced class weights, and all other hyperparameters were set to their default values.

In the “full” training mode, for each embedding, genes absent from that embedding were omitted from training in the corresponding fold. To isolate differences due to embedding quality rather than gene coverage, all methods were evaluated using the same holdout genes derived from the common intersection set.

To ensure consistent resources for assessing computation time, all evaluations were executed using 10 threads.

To compare methods across embedding data types, we first aggregated method-level AUROC scores by calculating the median performance for each data type group within each annotation set. Paired Wilcoxon signed-rank tests were then used to compare the median AUROC between groups.

### Paired gene interaction benchmark

For the SL and NG (both human and yeast) tasks, positive examples consisted of reported gene pairs for the respective genetic interaction. Negative examples were generated by permuting the genes present in the positive set to form unreported pairs and were randomly sampled to be 10 times larger than the number of positive examples. For TF target prediction, we included TFs with 500-1,000 targets (116 TFs total) and randomly downsampled to include 10,000 positive pairs. Negative examples were generated by permutation as with the SL and NG tasks.

For each embedding **e***_i_*, we constructed feature vectors for each gene pair (*i, j*) using three distinct combination strategies: element-wise summation (**f**_sum_ = **e***_i_* + **e***_j_*), the Hadamard (element-wise) product, (**f**_prod_ = **e***_i_* ⊙ **e***_j_*), and concatenation, (**f**_concat_ = [**e***_i_*; **e***_j_*]). Each combined feature was used as the input to an SVM classifier trained to predict gene-gene relationships.

Model evaluation followed a nested cross-validation scheme with 3-outer and 3-inner folds. More specifically, genes were partitioned into three disjoint outer folds; in each outer iteration, the outer test set included all gene pairs where both genes were only in the held-out outer fold. The remaining genes formed the outer training pool. The inner folds were used to tune the SVM C parameter within [0.1, 1, 10, 100, 1000] based on mean AUROC. The optimal C was then used to retrain the model on the full outer training pool and evaluated on the outer test set. All models used balanced class weights and a radial basis function kernel with otherwise default scikit-learn hyperparameters. All computations were performed using 10 threads for consistent timing. Gene pairs missing from an embedding in a given fold (in the full-gene variation of the task) were omitted.

### Gene set comparison benchmark

To evaluate how well gene embeddings capture relationships between sets of genes, we used our previously published ANDES [37] tool to assess relationships between: (1) matched pathways from GO and Kyoto Encyclopedia of Genes and Genomes (KEGG) [49], (2) disease and tissues using disease-associated genes (OMIM) and tissue-specific genes obtained from the Brenda Tissue Ontology [50], and (3) a temporal GO holdout.

For the matched GO-KEGG comparison, the GO gene sets were the same as in the gene-level prediction task and the KEGG gene sets were obtained from ConsensusPathDB [51]. We then obtained KEGG-GO mappings using the external database cross-reference to GO biological processes from the KEGG web portal for 52 KEGG pathways. To evaluate the capacity to capture functional similarity beyond counting overlapping genes, we excluded all overlapping genes between the KEGG– and GO-matched gene sets from the GO gene sets during the evaluation of each method.

In the disease-tissue analysis, we compared disease-associated gene sets with tissue-specific gene annotations. The OMIM gene sets were the same as the previous gene-level prediction task. The BRENDA annotations were obtained from TISSUE [52], and we included only annotations with experimentally determined annotations with a confidence z-score*>*1.64. To ensure tissue-specificity, we defined an “indirectly related set” for each term as tissue terms that cannot be reached without traversing through the root. Gene annotations were retained only if they were associated with fewer than 5% of the terms within this indirectly related set. Disease-tissue associations signify disease phenotypes observed in corresponding tissues and were made through manual curation of disease descriptions, resulting in 44 tissues terms associated with 79 disease terms. As the terms describe differing biology, unlike the GO-KEGG comparison, overlapping genes were retained when running ANDES.

For the temporal holdout task, we used the 19 GO terms rom the gene-level temporal benchmark. Each term’s annotations were divided into those annotated before 2024 and those annotated from 2024 onward (that had not been previously annotated). ANDES was then run to compare each pre– and post-2024 cohort.

### Embedding comparison

To compare the similarity between embeddings, we applied canonical correlation analysis (CCA) to each pair of embeddings. Because CCA can be computationally expensive in high-dimensional spaces, we first reduced each embedding matrix to its top 10 components using principal component analysis (PCA). These PCA-reduced embeddings were used solely for this comparison and not in any other downstream analysis. CCA identifies linear projections of two datasets that are maximally correlated with each other. Given two embeddings *X, Y* ∈ R*^N^*^×10^, CCA finds projection vectors *a* and *b* that maximize the correlation *ρ* = corr(*Xa, Y b*), computed as:

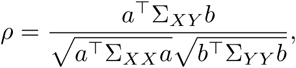

where Σ*_XX_,* Σ*_Y Y_*, and Σ*_XY_* are the covariance matrices within and between the embeddings. We then computed the coefficient of determination *R*^2^ to measure how well the canonical variables of one embedding predict those of the other. Formally, this is defined as 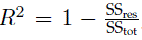, where SS_res_ is the residual sum of squares and SS_tot_ is the total sum of squares. An *R*^2^ of 1.0 indicates perfect linear predictability, while values near or below 0.0 indicate poor or worse-than-baseline performance. We calculated these pairwise *R*^2^ values for each pair of embeddings to quantify how much of the structure in one embedding is linearly predictable from another. All PCA and CCA computations were performed using the scikit-learn Python package.

To assess how embedding attributes contributed to predictive performance, we also evaluated the influence of different embedding attributes (algorithm, data input, and dimension size) to its average AUROC performance across each task. To do this, we fit an ordinary least squares regression with AUROC as the response variable against these three factors and performed a type II ANOVA across all benchmarks using the statsmodels Python library [53]. Each factor’s contribution was expressed as its proportion of the total sum of squares 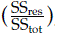, providing a normalized measure of relative influence across all benchmarks.

## Data availability

The embedding methods benchmarked are available through Zenodo at https://zenodo.org/records/16764517. Additional benchmark datasets are provided in the GitHub repository.

## Code availability

All code associated with this study is publicly available on GitHub at https://github.com/ylaboratory/gene-embedding-benchmarks under the BSD 3-clause open-source license.

## Acknowledgments

The authors thank Neel Mallipeddi and the members of ylaboratory for their feedback and suggestions. This work was supported by the Cancer Prevention & Research Institute of Texas (CPRIT RR190065) and the National Science Foundation (NSF DBI-2144534). VY is a CPRIT Scholar in Cancer Research.

## Supplementary Materials

**Table S1.** Average run times for training and testing all items in the benchmarking suite. Times are reported in seconds.

**Table S2.** ANOVA table comparing performance and embedding construction features.

**Figure S1.**
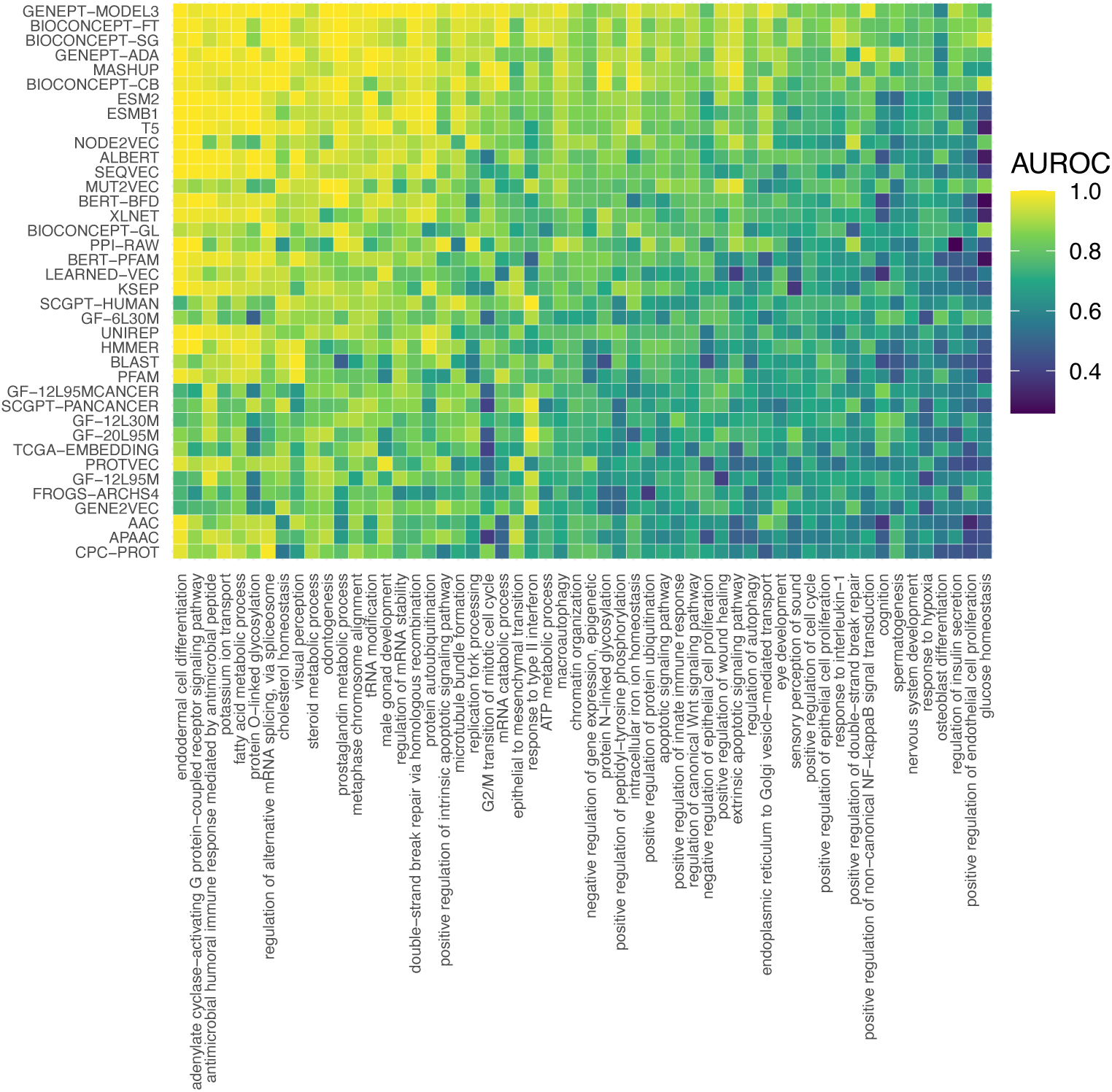
Single gene classification performance for each GO term. Heatmap showing the AUROC values for each embedding method and GO term.

**Figure S2.**
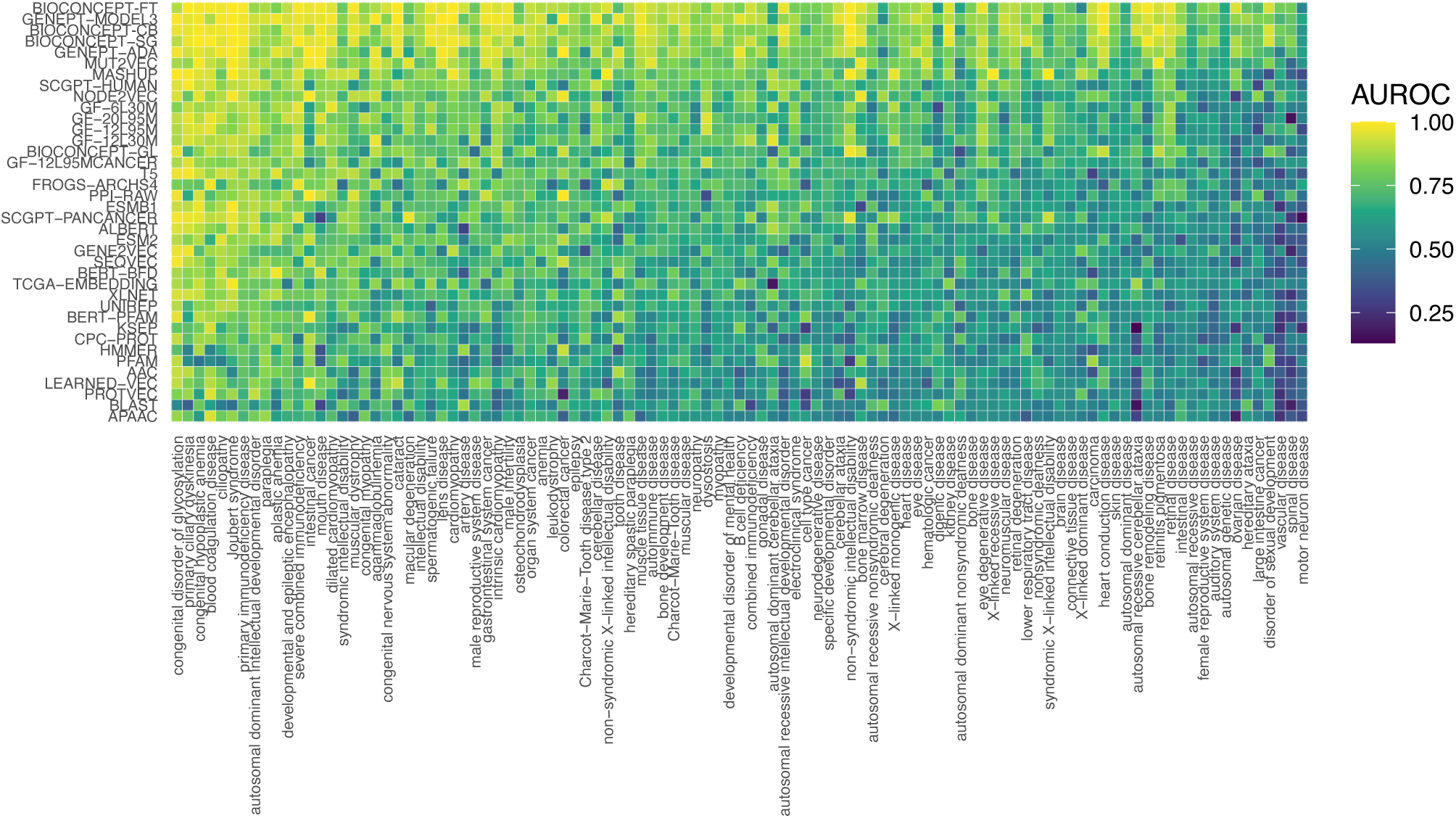
Single gene classification performance for each OMIM disease term. Heatmap showing the AUROC values for each embedding method and OMIM term.

**Figure S3.**
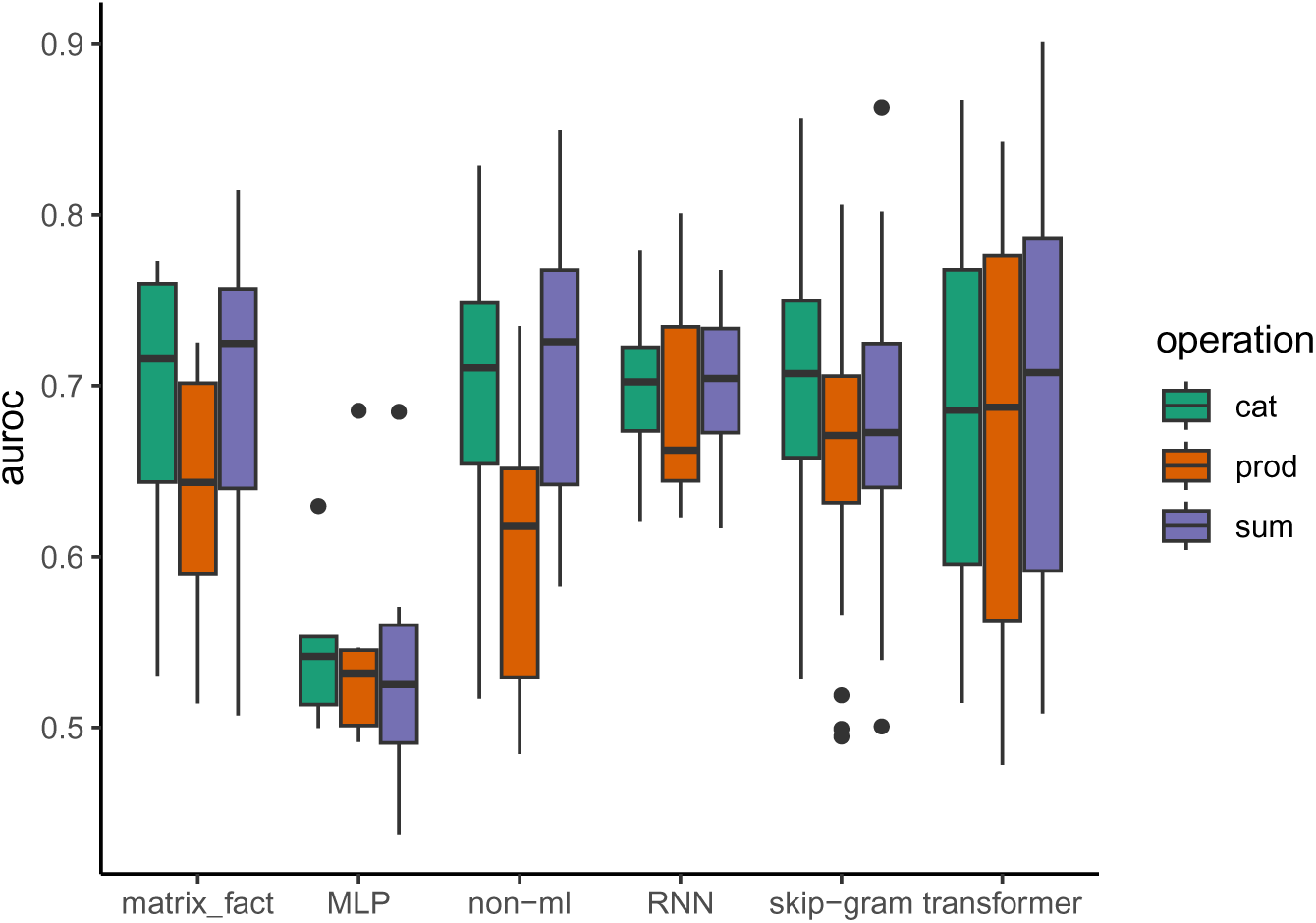
Paired gene performance as a function of embedding join operation. Boxplots detailing AUROC across the three paired gene tasks, SL, NG, and TF prediction as a function of their embedding joining operation.

**Figure S4.**
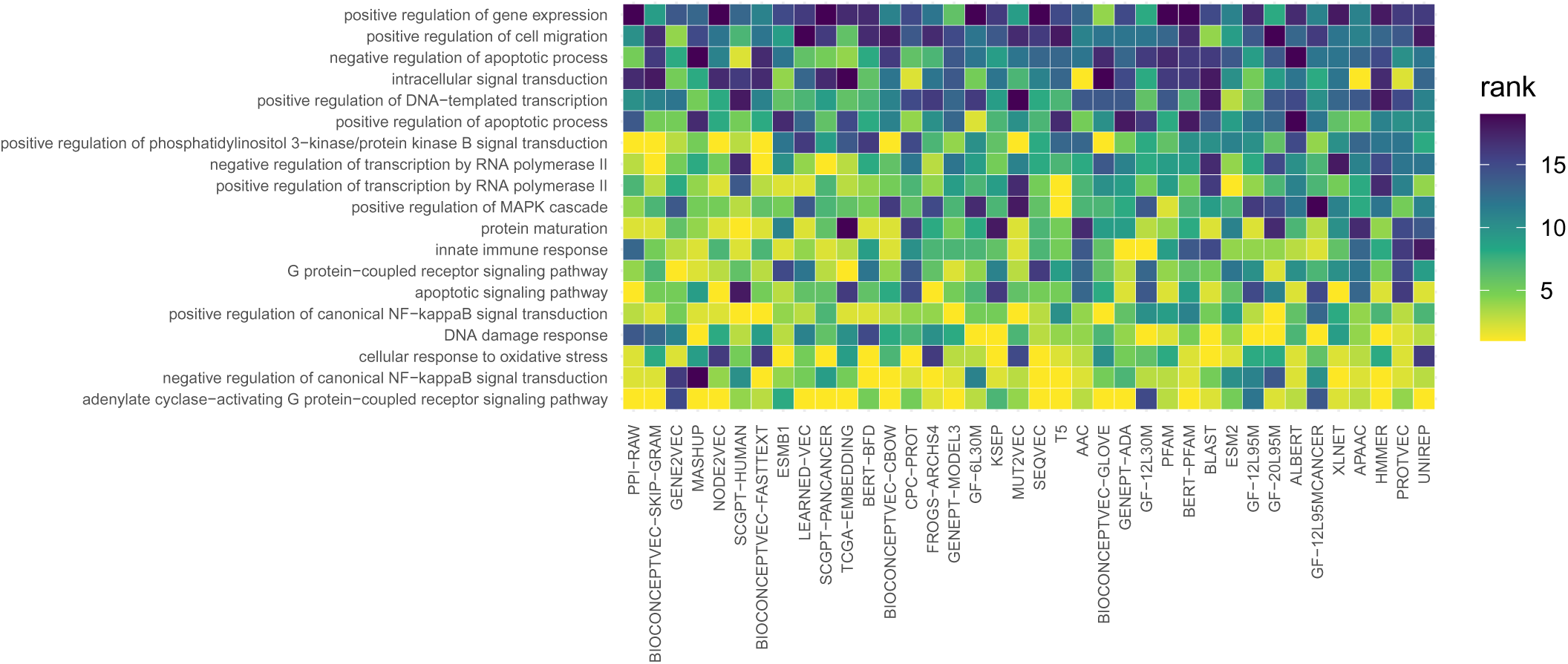
Gene set matching results for GO temporal holdout. ANDES ranks for each GO term with itself pre and post the 2024 holdout partition. Lower ranks indicate a better match between old and new terms.

**Figure S5.**
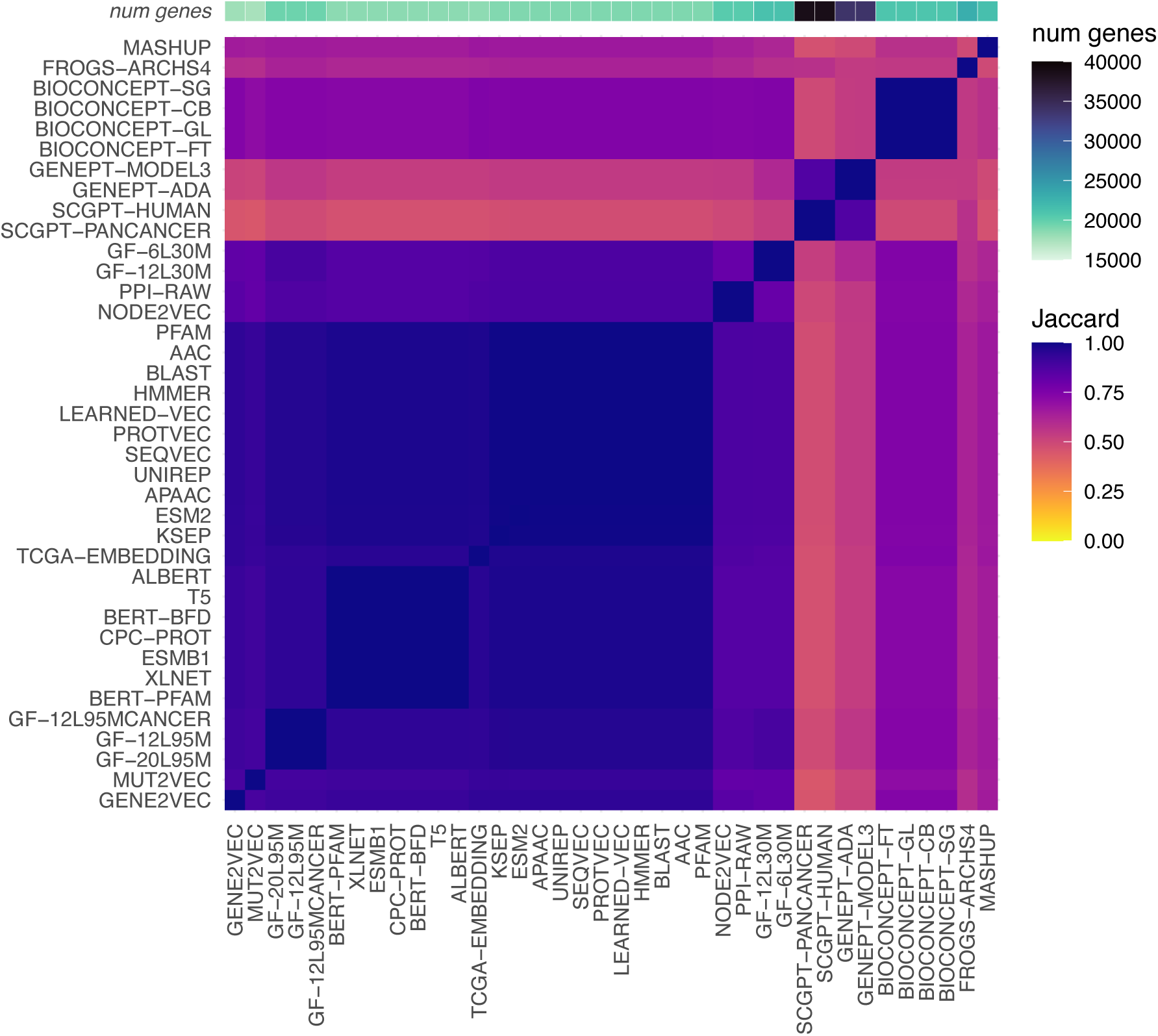
Embedding gene coverage. Heatmap showing the Jaccard overlap between included genes in each embedding.

